# Population Bursts in a Modular Neural Network as a Mechanism for Synchronized Activity in KNDy Neurons

**DOI:** 10.1101/2024.01.12.575324

**Authors:** Wilfredo Blanco, Joel Tabak, Richard Bertram

**Affiliations:** Department of Computer Science, State University of Rio Grande do Norte, Natal, Brazil; Department of Clinical and Biomedical Sciences, University of Exeter Medical School, Exeter, United Kingdom; Department of Mathematics and Programs in Molecular Biophysics and Neuroscience, Florida State University, Tallahassee, Florida, USA; Bioinformatics Multidisciplinary Environment of the Digital Metropolis Institute, Federal University of Rio Grande do Norte, Natal, Brazil

## Abstract

The pulsatile activity of gonadotropin-releasing hormone neurons (GnRH neurons) is a key factor in the regulation of reproductive hormones. This pulsatility is orchestrated by a network of neurons that release the neurotransmitters kisspeptin, neurokinin B, and dynorphin (KNDy neurons), and produce episodic bursts of activity driving the GnRH neurons. We show in this computational study that the features of coordinated KNDy neuron activity can be explained by a neural network in which connectivity among neurons is modular. That is, a network structure consisting of clusters of highly-connected neurons with sparse coupling among the clusters. This modular structure, with distinct parameters for intracluster and intercluster coupling, also yields predictions for the differential effects of the co-released transmitters neurokinin B and dynorphin. In particular, it provides one possible explanation for how the excitatory neurotransmitter neurokinin B and the inhibitory neurotransmitter dynorphin can both increase the degree of synchronization among KNDy neurons.

**Author summary:** Since the discovery of a small population of hypothalamic neurons that secrete kisspeptin, neurokinin B, and dynorphin (KNDy neurons), there has been interest in their role as a pacemaker for the pulsatile release of key reproductive hormones. A fundamental question is what mechanism coordinates KNDy neuron activity to generate population bursts. Optical imaging of the KNDy network at single-neuron resolution has revealed that individual KNDy neurons participate in many, but not all, population bursts. It has also shown that the order in which the neurons are recruited in each burst could be highly determined in some animals but not in others. We demonstrate here that these observations can be explained by a neural network with a modular structure. We also show how such a structure can explain the paradoxical finding that both the excitatory neurotransmitter neurokinin B and the inhibitory neurotransmitter dynorphin can act to increase the degree of synchronization among the KNDy neurons.

## Introduction

The gonadotropins luteinizing hormone (LH) and follicle-stimulating hormone (FSH) play key roles in fertility through their actions on other hormones and gamete development [1]. The gonadotropins are secreted by gonadotrophs located in the anterior portion of the pituitary gland. Secretion of these key reproductive hormones is controlled by hypothalamic gonadotropin-releasing hormone (GnRH) neurons [2, 3], which release GnRH into the hypophyseal portal bloodstream in males and females in pulses [4, 5], driven by bursts of electrical activity [6, 7]. GnRH must be delivered in a pulsatile manner since continuous delivery desensitizes gonadotropin release [8, 9]. The mechanism for the synchronous release of GnRH from the GnRH neurons has been a matter of investigation for many years, pushed forward by the discovery in 2003 that mutations in the gene encoding the G protein-coupled receptor for kisspeptin led to hypogonadotropic hypogonadism [10, 11]. We now know that pulsatile GnRH activity is coordinated by a small population of kisspeptin (Kiss)-releasing neurons in the arcuate nucleus of the hypothalamus that also release neurokinin B (NKB), and dynorphin (Dyn), and are known as KNDy neurons [12–18]. In short, the pulsatility of GnRH neuron activity reflects pulsatility in KNDy neuron activity, with kisspeptin serving as the output of the KNDy network to the GnRH neurons [17]. The obvious next question is what mediates the synchronous episodic activity in the population of KNDy neurons? These neurons are interconnected, and the “KNDy hypothesis” suggests that release of the stimulatory neurotransmitter NKB from KNDy neurons to neighboring KNDy neurons starts an episode of electrical activity, while a delayed action by Dyn terminates an episode [19–21]. There is substantial evidence supporting this hypothesis, reviewed in [17, 19], but recent data supports an alternate hypothesis in which coupling among KNDy neurons through glutamate is the essential ingredient for the coordinated rhythmic activity of the neurons [22, 23]. According to this hypothesis, glutamate provides the excitation responsible for initiating each episode of electrical activity through actions on AMPA receptors, while either synaptic depression or the buildup of intracellular Ca^2+^ acting on Ca^2+^-activated K^+^ channels within the cells ends each episode. NKB and Dyn then serve as modulators of the rhythmic activity of the population of neurons [22], with NKB being particularly important in brain slice studies from female mice [23]. In addition to receptors for NKB and Dyn, KNDy neurons have been shown to express AMPA receptors [24], and to release glutamate onto KNDy neurons [25], which are essential elements of this “glutamate hypothesis”.

The goal of this article is not to weigh in on the validity of either the KNDy or glutamate hypothesis. Instead, through mathematical modeling, it aims to demonstrate how an implementation of the glutamate hypothesis along with a modular network structure can account for experimental findings reported in two recent studies [22, 26]. In so doing, we also show how apparent discrepancies in some results of these studies can be explained by heterogeneity in the modular network. The experimental findings were obtained using GCaMP transfection to measure Ca^2+^ fluorescence in individual KNDy neurons either *in vivo* [22, 26] or in brain slices containing a portion of the arcuate nucleus [22, 23]. With these measurements, it was possible to examine the activity of many KNDy neurons simultaneously. The findings we wish to explain with the model are the following:

1. Why do many neurons participate in some episodes of activity, called “synchronization events” (SEs), but not all [22, 26]?
2. Are there “leader cells” that consistently fire first during SEs [26], or is the temporal order more random [22]?
3. How can the actions of the excitatory modulator NKB and the inhibitory modulator Dyn both lead to a greater degree of synchronization during SEs [22]?

Modular networks are characterized by clusters of highly-connected nodes with sparse coupling among the clusters. In the context of neural networks in which coupling is through excitatory glutamatergic synapses, this structure leads to a high degree of coordinated activity among cells in a cluster, and some weaker coordination among the clusters. This structure gives rise to two qualitatively different types of connections: the intracluster connections (intraCC) and the intercluster connections (interCC). While the two are implemented in the same way in the model, we show that their roles on the synchronisation behavior of the population of neurons are different. This characteristic of modular networks provides simple answers to the three questions above.

## Methods

We do not assume any special properties (such as rhythmic bursting) for the KNDy neurons, so we model them using a reduced Hodgkin-Huxley model, as we used in previous studies [27–30]. In Ca^2+^ measurements from intact animals, SEs occur once every 5-20 min [15, 22, 26]. Since the time scale for electrical impulses is in milliseconds, replicating the long intervals between SEs would require very long computations. Our focus is on the impact of a modular network structure, and not on accurately reproducing the time scale of KNDy network behavior, so we simulate SEs with a much smaller inter-SE interval of approximately 1 s. We first describe the single-cell model, then the way that the network is implemented. All parameter values are given in Table 1.

**Table 1.** Parameters of the network model.

| Parameter | Description | Value |
| --- | --- | --- |
| $g_l$ | Leak conductance | 0.1 mS/cm<sup>2</sup> |
| $V_l$ | Leak reversal potential | -49.4 mV |
| $\bar{g}_{Na}$ | Sodium conductance | 36 mS/cm<sup>2</sup> |
| $V_{Na}$ | Sodium reversal potential | 55 mV |
| $\bar{g}_k$ | Potassium conductance | 12 mS/cm<sup>2</sup> |
| $V_k$ | Potassium reversal potential | -72 mV |
| $\bar{g}_{syn}$ | Max. synaptic conductance | 3.6 mS/cm<sup>2</sup> |
| $V_{exc}$ | Excitatory reversal potential | 10 mV |
| $I_{bkg}$ | Constant background current | -10 to 5 μA/cm<sup>2</sup> |
| $\alpha_a$ | Synaptic activation rate | 1 ms<sup>-1</sup> |
| $\beta_a$ | Synaptic decay rate | 0.1 ms<sup>-1</sup> |
| $\alpha_s$ | Synaptic recovery rate | 0.0015 ms<sup>-1</sup> |
| $\beta_s$ | Synaptic depression rate | 0.12 ms<sup>-1</sup> |
| $V_{th}$ | Threshold for activation | -20 mV |

Parameter values used in model simulations.

### The single-cell model

The intrinsic behavior of cell *j* is described by two differential equations, one for the cell’s membrane potential (*V*_*j*_) and one for the activation variable of a delayed rectifying K^+^ current (*n*_*j*_):

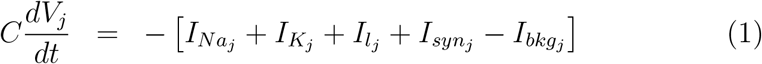

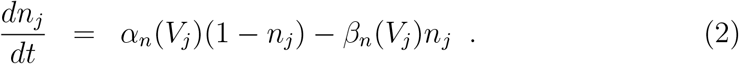

The Na^+^ current is simplified to assume instantaneous activation and utilizes the almost-linear relationship between its inactivation variable and the activation variable for K^+^, as described in [31]:

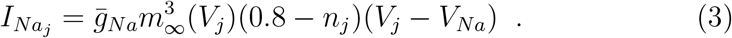

The K^+^ and leak currents are, respectively:

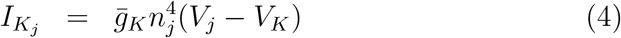

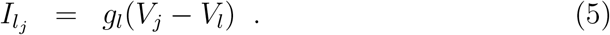

Each model neuron receives excitatory synaptic input from one or more other neurons:

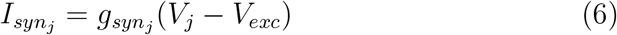

where *V*_*exc*_ is the excitatory reversal potential. The synaptic conductance is the sum of input from all neurons innervating neuron *j*. Finally, the synaptic conductance onto neuron *j* is

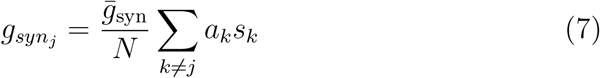

where the summation is over all neurons innervating neuron *j*, 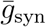 is the synaptic conductance strength parameter, and *N* is the total number of neurons in a cluster (*N* = 50). Each of these neurons has an activity level, *a*_*k*_ ∈ [0, 1], and a synaptic efficacy *s*_*k*_ ∈ [0, 1]. The activity level increases with each presynaptic spike and represents the “synaptic drive” from neuron k to other neurons. The synaptic efficacy reflects synaptic depression, so it declines with frequent presynaptic activity. The differential equations are:

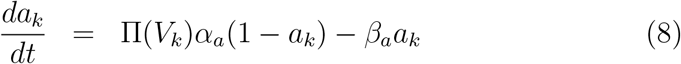

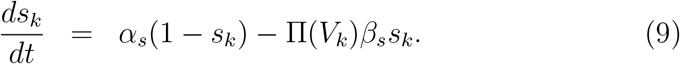

The increasing sigmoidal function 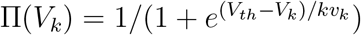 reflects the synaptic release process that occurs when the presynaptic voltage *V*_*k*_ goes past a threshold *V*_*th*_ during an action potential. When this happens, *Π*(*V*_*k*_) increases from a value ≈ 0 to a value ≈ 1 for a short period of time, before returning to ≈ 0. The *α* and *β* parameters are rate constants.

The final current in the voltage equation is a constant background current, *I*_*bkg*_, that sets the excitability of the cell. For each cell, this is drawn once randomly from a uniform distribution over the range -10 to 5 *µ*A/cm^2^, ensuring the heterogeneous activity of the network (on average, 10% of the cells spike in the absence of synaptic input).

### The modular network

For all simulations, we use a population of 250 model neurons. We form 5 cell clusters of 50 neurons with a high degree of interconnectivity within each cluster (Fig 1A). Cells within these clusters are connected to cells in other clusters with many fewer links (Fig 1B). To generate a cluster, we first set the coupling parameter for the fraction of cells within the cluster that a neuron should connect to (called “intraCC” for intracluster coupling). The same value is used for all 5 clusters. Then pseudo-random numbers are generated to determine which connections are actually made. A similar process is done for intercluster coupling. The coupling parameter “interCC” is then the fraction of all possible connections between clusters that are actually made.The extent of coupling within each cluster is large in our model, but the coupling strength of each connection is small, so that stimulation of any one neuron so that it fires tonically is typically insufficient to evoke firing in neurons that it synapses onto. This is consistent with the brain slice experiments of Han et al., in which they found that stimulating one neuron rarely had an effect on the behavior of the other neurons that they were examining [22].

**Fig 1.**
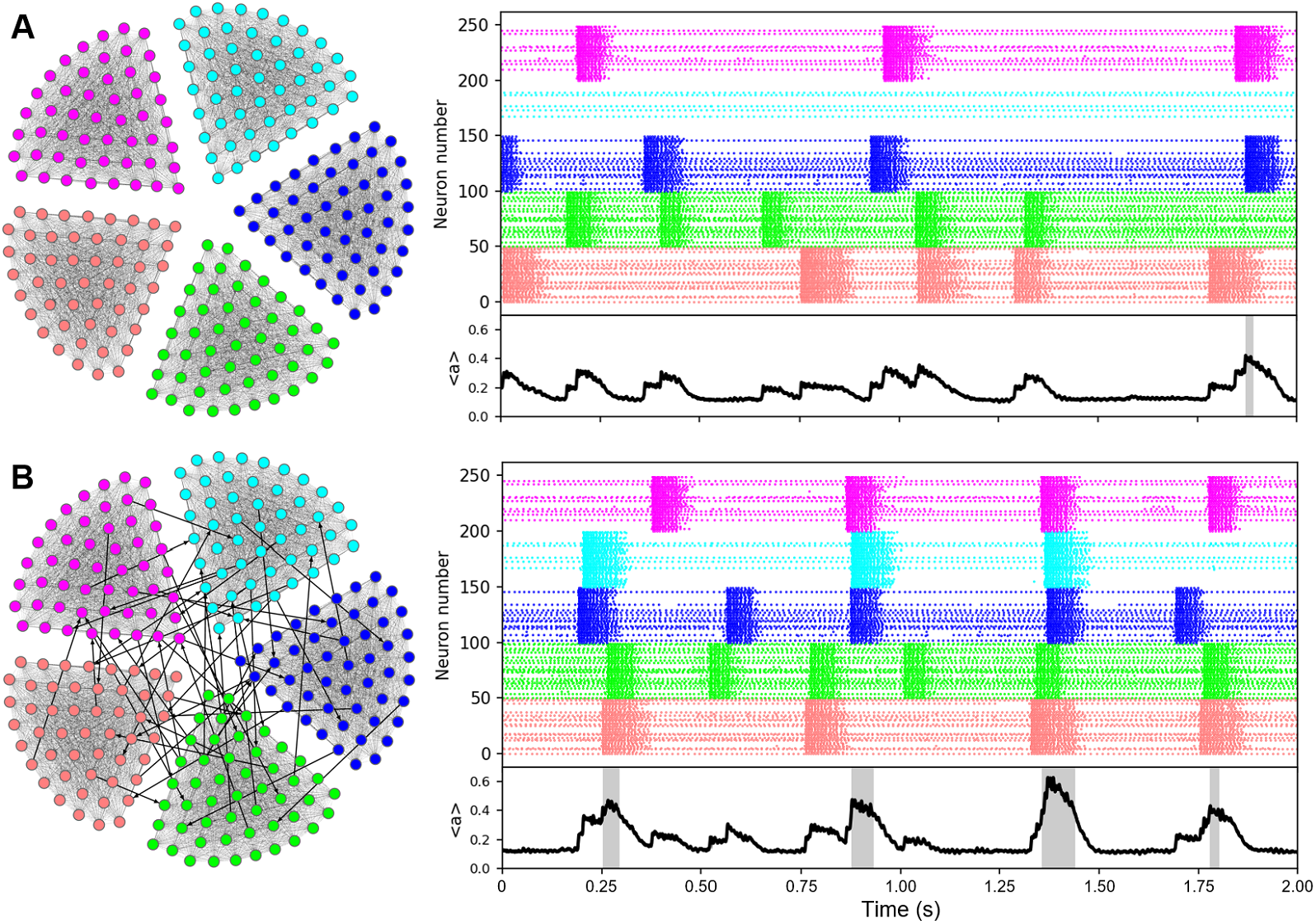
Cluster events and synchronization events in the modular network. The model neural network has 5 highly connected cell clusters (left). Each of these is capable of producing population bursts of activity or “cluster events” (CE), as seen in the raster plots (right), color-coded to indicate the cluster that the cells are part of. The bottom black curve over time is the activity averaged ⟨*a*⟩ over all 250 cells in the population. When overlapping events occur in 3 or more clusters, ⟨*a*⟩ is above the threshold for what we refer to as a synchronization event (illustrated with gray shading) A: Without interconnections among the clusters (intraCC=100%, interCC=0%). B: Interconnections among the clusters, though sparse, can synchronize the cluster events. We have sped up the synaptic efficacy variable to decrease the simulation time scale for producing synchronized events from hour to second. (intraCC=100%, interCC=0.1%)

The network activity (denoted as ⟨*a*⟩) is calculated by averaging over all cell activities *a*_*k*_ (Fig 1, black traces). When all clusters fire together, ⟨*a*⟩ rises to ≈ 0.6, and we define a synchronization event to occur when 3 of the 5 clusters are simultaneously active, so when ⟨*a*⟩ ≥ (0.6)(0.6) = 0.36 (Fig 1, gray shading).

The model and simulations were implemented using the Eclipse IDE for C and C++ with the MinGW gcc v12.2.0 compiler. Ordinary Differential Equations (ODEs) were solved using the Runge-Kutta fourth-order (RK4) method with time step 0.01 ms. The output .txt files were processed using Python (v3.10.9) and Matplotlib (v3.7.0) to generate the figures. The graph in Figure 1 was created using Gephi [32]. Source code can be downloaded from www.math.fsu.edu/bertram/papers/neuron.

## Results

### The modular network exhibits a mix of partially and fully synchronized events

The model modular network consists of 5 cell clusters with extensive intracluster coupling (Fig 1A left) quantified by the coupling percentage “intraCC”, and much less extensive intercluster coupling (Fig 1B left) quantified by the coupling percentage “interCC”. Each cluster contains 50 neurons described by Hodgkin-Huxley-like models. Although each cluster has the same intraCC, the intracluster network structure is determined randomly, so will differ from cluster to cluster. In addition, the background currents in the model neurons are determined randomly from a uniform distribution (see Methods), so neurons in the network have different excitability. For these reasons, some clusters are more active than others. The raster plots in Fig 1A, where there is no intercluster coupling, illustrate this. The top (pink) cluster has cells that are tonically active, and those that are inactive. However, there are three instances when the entire cluster is active, i.e., there is a population burst that we refer to as a “cluster event” (CE). The onset is triggered by a few initiating cells that spike and, due to the extensive coupling, cause others in the cluster to spike [28]. An episode terminates due to the buildup of synaptic depression that reduces the coupling between the cells to a level that is eventually too low to continue the regenerative activity [28, 33].

The second (cyan) cluster exhibits a quite different activity pattern. Although there are some tonically active cells, at no point during the 2 s simulation is there a CE. The fourth (green) and fifth (red) clusters, on the other hand, exhibit 5 CEs over the 2 s simulation time. Clearly then, with the randomness in the coupling and distribution of background currents, the cluster activity is very heterogeneous.

The bottom panel shows the activity variable, *a*_*k*_, averaged over the population of 250 cells, denoted as ⟨*a*⟩. The time course of ⟨*a*⟩ shows the timing of bursts within a cluster, and when bursts occur in two or more clusters simultaneously this is reflected in the amplitude of the ⟨*a*⟩ deflection. Thus, ⟨*a*⟩ can be used as a metric to determine whether the bursts in the clusters are synchronized. There were several occasions in which bursts in two clusters overlapped in Fig 1A, even though the clusters are not coupled, and this is reflected in larger deflections in ⟨*a*⟩.

When a low level of intercellular coupling is added (interCC=0.1%) there are, not surprisingly, more instances of coordinated bursting. In the 2 s simulation shown in Fig 1B, there are instances of coordinated bursting in 3, 4, or all 5 clusters. When three or more clusters exhibit overlapping bursting we call this a “synchronization event” (SE), since the majority of cells in the population are spiking simultaneously. These SEs are highlighted in gray in the time course for ⟨*a*⟩. In this case, a cell is receiving, on average, 0.001*200 = 0.2 synapses from other clusters than the one it belongs to, so a cluster receives about 10 connections from the four other clusters – this is enough to produce synchronized cluster events.

An observation made in [22, 26] in Ca^2+^ imaging studies of KNDy neurons *in vivo* is that many of the neurons participated in some, but not all, of the synchronization events. We examine this in the modular network in Fig 2, which shows activity traces (the variable *a*) for 3 randomly-chosen neurons from each of the 5 clusters. In this network, there is complete coupling within each cluster (intraCC=100%) and weak coupling among clusters (interCC=0.4%). Over a period of 5 sec, this network produces 9 SEs. All 15 cells participated in the first SE. However, only 12 participated in the fifth SE; cells in the blue cluster did not exhibit a CE, and so remained inactive (indicated by a red X). In the seventh and eighth SEs, different clusters of cells did not participate, from the magenta and cyan clusters. An interesting case is SE 6, where cells in the green cluster had a burst immediately before, but not during, the SE. These likely contributed to the SE initiation. Also, one of the cells in the cyan cluster was active during the SE (orange circle), while the other two from the same cluster were inactive. This illustrates that cells sometimes participate in SEs even though their cluster does not produce a CE. Overall, the figure shows that many cells in the modular network participate in some, but not all, of the SEs, as reported in [22, 26].

**Fig 2.**
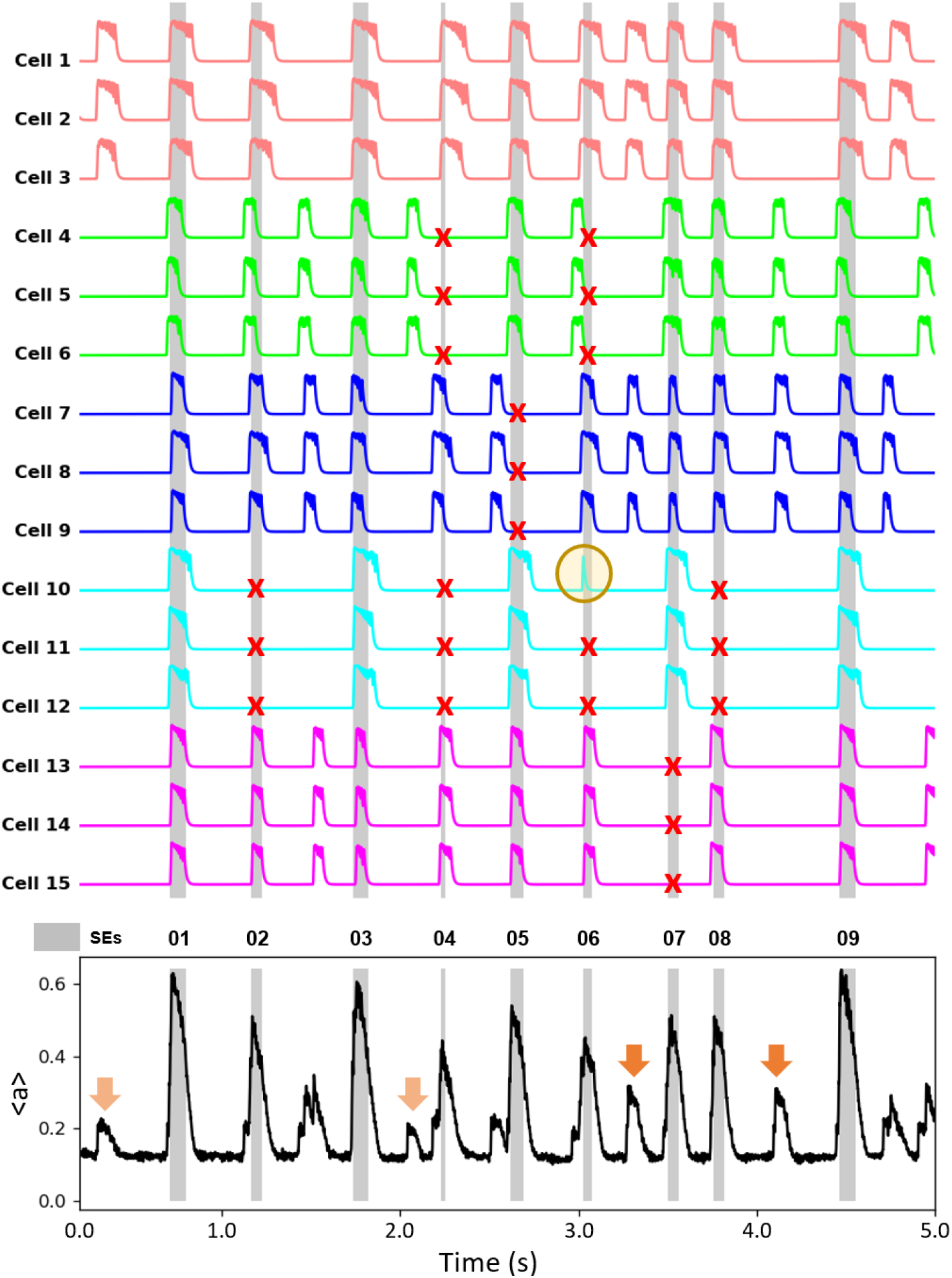
Synchronization events and miniature synchronization events in the modular network. The activity time courses (the variable *a*) for 3 model neurons selected randomly from each of the 5 clusters. Most cells participate in some, but not all, of the synchronization events. A red X highlights an instance in which a cell did not participate, and an orange circle highlights an instance in which a cell did participate, but other cells in its cluster did not. The bottom panel shows the average activity across the network, with SEs indicated by gray bars. Arrows in the bottom panel highlight a few (but not all) instances of mini-SEs, where one cluster (light orange arrow) or two clusters (dark orange arrow) produced bursts of activity. IntraCC=100%, interCC=0.4%.

Another phenomenon shown in Fig 2 is that the averaged activity of the population has a mix of large increases, the SEs, and smaller ones (marked with orange arrows). These smaller events reflect burst activity of only one or two clusters. Smaller events in the population activity were referred to as “miniature synchronization events” in [22], and we use this nomenclature for the smaller events that occur in the modular network.

### Leader cells are possible, but not guaranteed

Are there leader cells that consistently fire near the beginning of SEs and therefore serve as triggers for the SEs? This question was addressed in vivo by both Moore et al. [26] and Han et al. [22]. The former found that there was indeed a set of KNDy neurons for which the Ca^2+^ level consistently reached its peak at the beginning of an SE in which it participated. Other KNDy neurons consistently fired in the middle of an SE, and others consistently fired at the end, indicating that they were recruited to fire by other neurons (Fig 4 of [26]). The latter study showed much more flexibility in the firing order of KNDy neurons. In some animals there appeared to be a similar preferred firing order of KNDy neurons as in [26], while in other animals there was no consistent temporal ordering (Fig 1 of [22]). Similar results were shown for SEs that occurred in vitro in brain slices (Fig 2 of [22]). How can these conflicting data be reconciled?

We examined this question in the modular network using different combinations of the connectivity parameters. Figure 3A shows one example with intraCC=55% and interCC=0.6%. We examined the activity of 15 neurons (3 from each cluster) as this is a typical number of neurons recorded simultaneously [22, 26]. Activity traces for the 15 neurons were analyzed, starting with time courses of the neurons during two SEs. From these, it is apparent that cells in the green cluster began firing first in the two SEs, while those in the blue or magenta cluster began last. The firing order for 8 SEs is shown in the left table. Each entry of this table gives the order that firing began, and the boxes with light coloring represent firing near the beginning of an SE and dark coloring represents firing late in the SE. From this, we see that the cells in the green cluster consistently began firing first, they are leader cells, while those in the blue and magenta clusters mostly began near the end of the SEs. To bring out the temporal ordering more clearly, the table was reorganized so that the ordering of the rows is based on the typical order when firing began during SEs; cells that typically began firing first are placed in the top row, and those that typically began last are placed in the bottom. In this rightmost table, the shading variation from top to bottom clearly demonstrates consistency in the temporal order of spiking during SEs. Finally, the data are shown as scatter plots, with the order in which spiking began on one axis and the cell ID on the other, with cells that typically began first placed on the bottom and those that typically began last placed on the top. The data are color-coded to correspond to the cluster that the corresponding neuron is part of. When organized this way (as was also done in [22] and [26]), the consistency of the temporal order of firing can be quantified with the Pearson correlation coefficient (R). The large value of the R (0.85) demonstrates that there is consistency in the temporal order in which spiking began during SEs in this example network. Also, with the color coding, it is clear that neurons in the green cluster consistently fired first, they are leader cells, while those in the magenta and blue clusters consistently fired last and so are follower cells. This ordering reflects the level of activity produced by the clusters without intercluster connections. That is, without any intercluster connectivity the neurons in the green cluster produce the most frequent bursts of activity, followed by neurons in the red cluster, with neurons in the magenta and cyan clusters not active at all without intercluster coupling.

**Fig 3.**
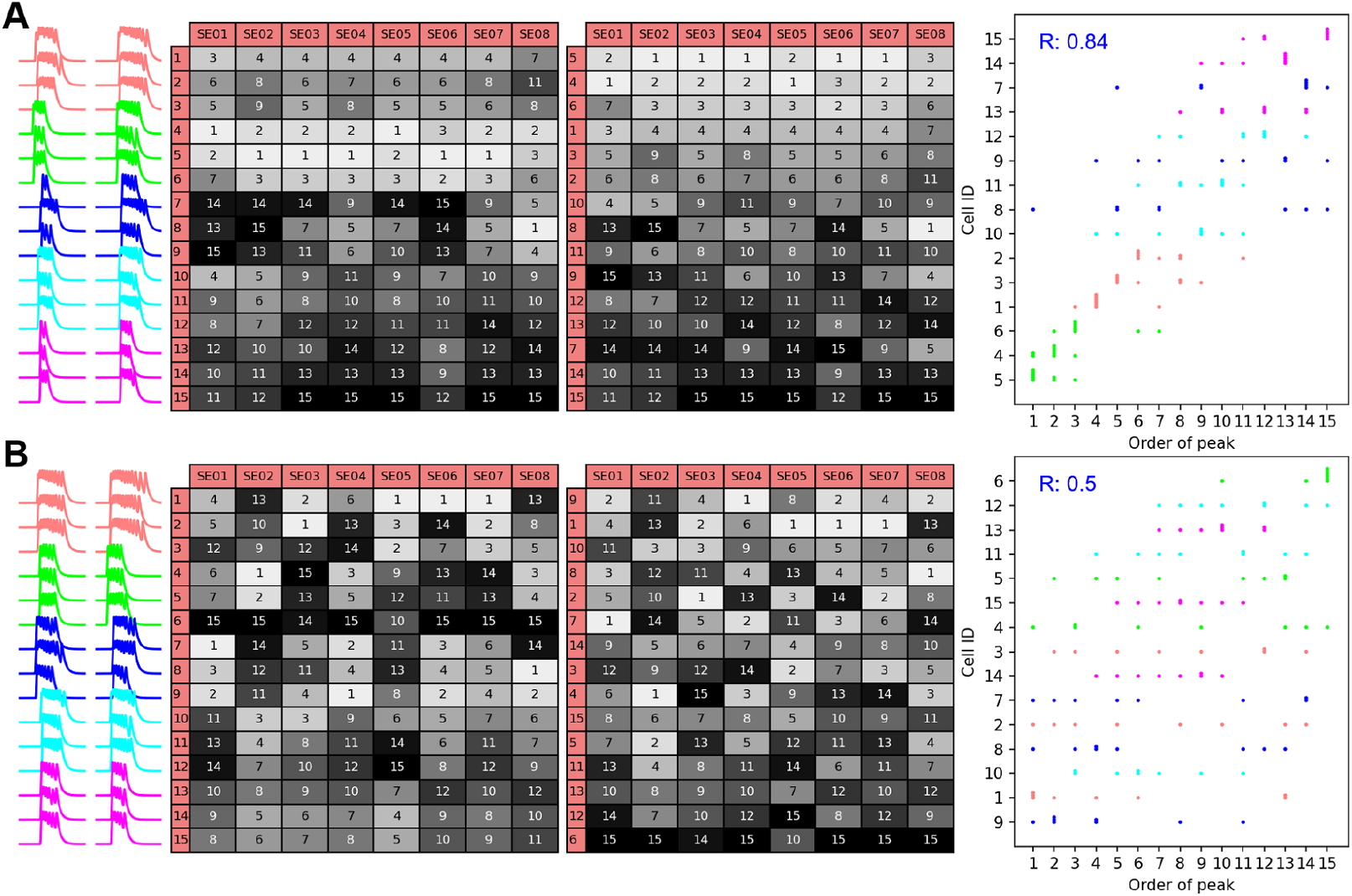
Network connectivity parameters determines whether there are leader cells. A: Activity time courses of 15 neurons selected from all 5 clusters during two SEs (left, SE01 and SE02). The left table shows the temporal order of firing of the 15 neurons in 8 SEs; light shading indicates that the cell fired early in an SE. The right table is a reorganization of the left table so that cells that typically fire early are placed in the top rows. The scatter plot indicates spike timing with those cells spiking early in the SEs placed on the bottom. There is a strong correlation (R = 0.85), indicating the presence of leader cells. The data points are color-coded according to the cluster that the corresponding neurons are part of. IntraCC=55%, interCC=0.6%. B: With a different value of the intracluster coupling parameter there is much less consistency in the temporal order of spiking during SEs (R = 0.5). IntraCC=75%, interCC=0.6%.

Another example network, with a larger value of intraCC, produced different results (Fig 3B). While neurons in the red cluster often began to fire early in an SE, they also sometimes began much later. Cells in the magenta cluster most often began to fire toward the end of an SE, but sometimes began at the start of an SE. The lack of consistency in the temporal order of spiking is seen most clearly in the scatter plot, where the R (0.5), is much lower than in the previous example. This value is very similar to what was reported in most of the data from [22]. In addition, it is evident from the color coding that there are no clusters that consistently fired first. This likely reflects the fact that when intraCC is increased all the clusters produce bursts of activity, even without intercluster coupling, making it more likely that any cluster can start an SE by recruiting other clusters to fire.

The last figure raises the question of whether consistent temporal firing is more likely with some network coupling parameters than others. To investigate this, we examined the temporal order R for a grid of coupling values (intraCC ∈ [50, 100]% and interCC ∈ [0.2, 2]%). The first heat map of Fig 4A reflects one simulation for each pair of coupling parameter values in the grid. It shows that the R is highest for small values of intraCC and large values of interCC. The second through fourth heat maps use the same grid of coupling parameter values, but the networks are different, since edges between neurons are chosen randomly in accordance with the coupling parameters. In each case, the temporal firing R is generally higher for small intraCC and large interCC values. This is also reflected in the mean of R values, averaged over 8 grids of simulations (the 4 shown, plus 4 others). If each pair of coupling parameters represents an individual animal, then depending on which animal is examined there will be great consistency in the temporal ordering of cell firing during SEs, with leader cells and follower cells, or the ordering will be much less regular.

**Fig 4.**
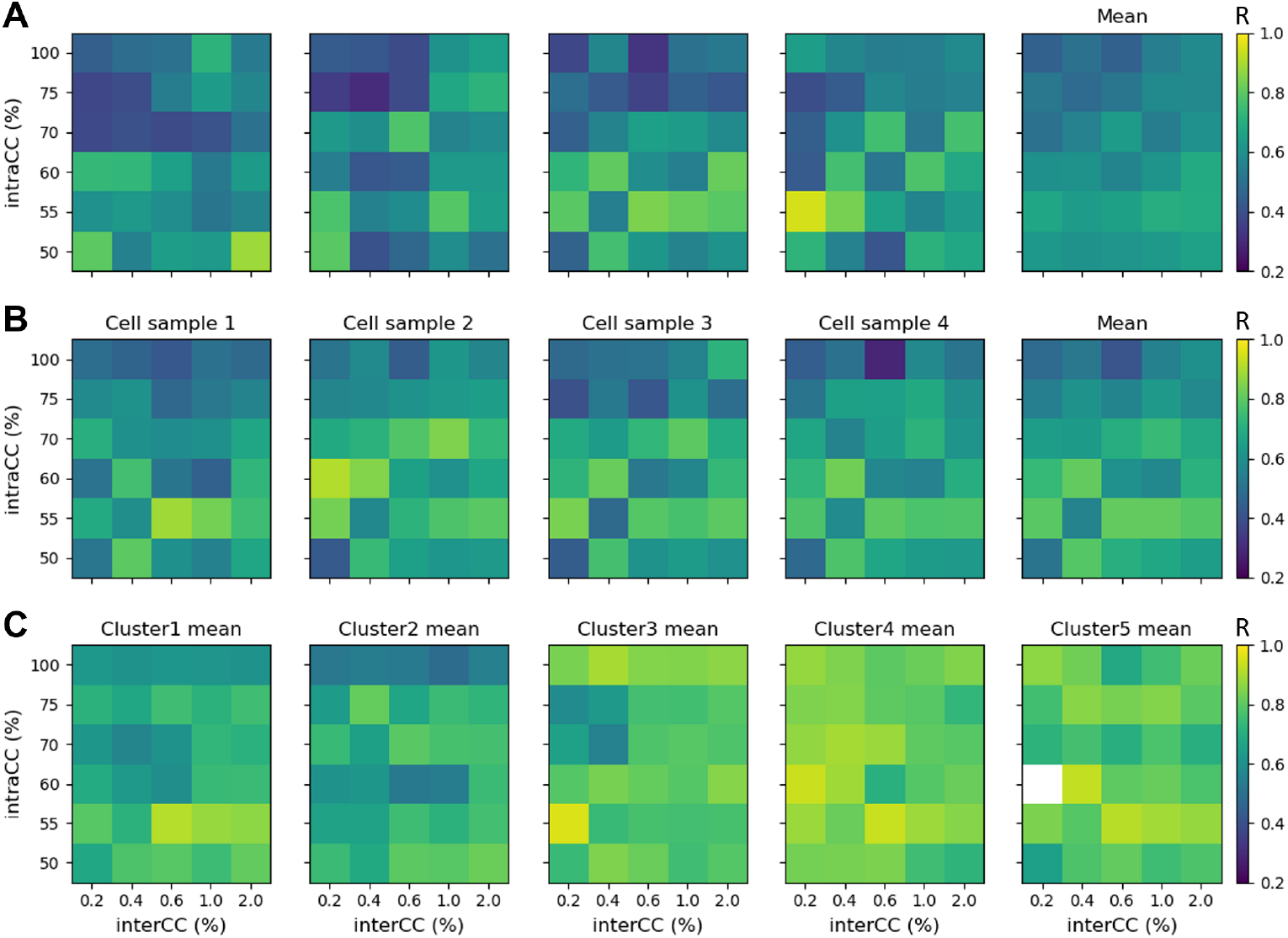
Some coupling parameter combinations are better than others for producing a consistent temporal ordering of cell activity during SEs. Each element of the heat map shows the temporal order correlation coefficient (R) value for a network with a specific pair of coupling parameters, intraCC and interCC. Light colors indicate high R. A: Four simulations of networks across the entire grid of coupling parameters. The calculation of R is based on the activity of a fixed set of 15 cells, with 3 chosen randomly from each cluster. The last heat map is the average of 8 grids of simulations (these 4, plus 4 others). The networks used in the third grid are used again for the next two panels below. B: The choice of the 15 neurons used in the calculation of the R is random across the network, and different random samplings are made in each of 4 simulations. The last heat map is the average across the 4 simulations. C: The choice of 15 neurons used in the calculation of R is made from a single-cell cluster. For each heat map, four random samplings of 15 neurons were made, and the R was computed. The heat map is the average of these. In one case (white element in the heat map) the cluster produced no activity bursts, so an R could not be computed.

In Fig 4B we explored how the 15 cells chosen for the calculation of the temporal order R affects the results. For each pair of coupling parameters chosen a network is constructed, and this same network is used for all simulations in panel B. For each simulation, a set of 15 neurons is chosen randomly from the network, and the temporal order R is determined. In the last heat map, the R values across the four simulations are averaged. Just as in panel A, the R is largest at lower values of intraCC and higher values of interCC. Thus, the selection of cells used in calculating the R is not important, as long as they are selected from all clusters of the modular network. Again, this suggests that consistent cell recruitment order reflects consistent cluster recruitment order.

What happens to the temporal order if all 15 neurons examined are selected from the same cluster? This is shown in Fig 4C, where each heat map corresponds to a selection from one of the five clusters (using the same networks that were constructed in panel B). In each simulation, 15 neurons were chosen randomly from a single cluster and the temporal order R was calculated. This was done four times, corresponding to selection from one of 5 different clusters, and the average of these is shown in the heat map. In all of these cases, the temporal order R is high, with little variation across coupling parameters. This demonstrates that a major source of the flexibility in temporal ordering seen in panels A and B is variation in the order of activity of the clusters, which differs from SE to SE. Thus, if laboratory measurements of cell activity only captured the activity of a single cluster, then a high degree of temporal order would be observed, with distinct leader and follower cells.

### Differential effects of changes in the coupling parameters

In [22], small “miniature SEs” (mSEs) were distinguished from SEs as being significantly smaller and therefore reflecting a smaller degree of synchronous neural activity. Following this nomenclature, in the simulations it is natural to categorize events in which a majority of the clusters (3 or more) fire together as SEs, and events in which one or two clusters fire together as mSEs. Hence, if we define NCE_*k*_ as the number of events in which *k* clusters were simultaneously active (i.e., in which there were *k* “cluster events”), then the number of mSEs throughout a simulation is NCE_1_+NCE_2_ and the number of SEs (which we refer to as NSE) is NSE=NCE_3_+NCE_4_+NCE_5_. Figure 5 A shows the number of cluster events for different values of the interCC parameter in simulations of 40 s duration. For example, with interCC=0.2% (top left histogram, with tan shading), 62 events were single-cluster events (NCE_1_), while all clusters fired together in only 22 events (NCE_5_). The number of synchronization events is NSE=39 (3+14+22). The top panel of Fig. 5B shows the average activity of each cluster during 5 s of the simulation corresponding to this same coupling parameter value. The number of CEs is shown on the right. Several CEs do not recruit all the other clusters into full-blown SEs.

**Fig 5.**
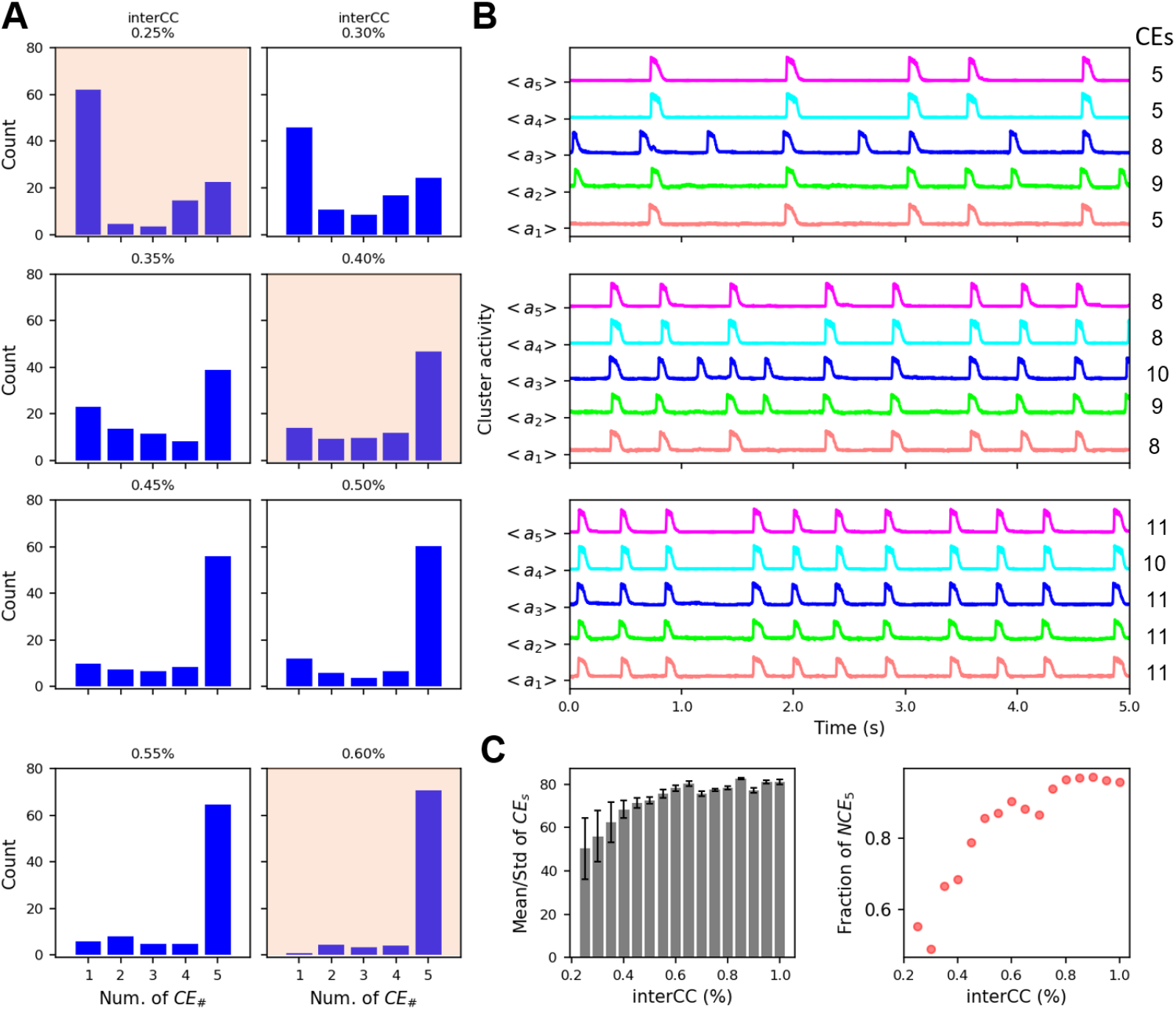
The number of cluster events and the degree of synchronization increases with an increase in the intercluster coupling. A: Histograms showing the number of events in which 1, 2, …, 5 clusters fired together during simulations with 40 s duration. The interCC is increased moving from top to bottom. B: Average activity time courses of the 5 clusters during 5 s of simulation time, corresponding to the histograms with tan shading in the previous panel. The number of CEs is shown on the right. C: (left) Mean number of CEs, along with standard deviation, for the range of interCC values explored in the histograms and over simulations with 40 s duration. (right) The fraction of synchronization events (events with 3 or more clusters active) in which all 5 clusters are active, NCE_5_/NSE. IntraCC=60%.

When the intercluster coupling is increased there is a clear shift in the histogram, so that by interCC=0.35% the vast majority of events are SEs, and most of these have all clusters firing in synchrony. With interCC=0.60% almost all CEs are in SEs with all clusters firing together. This increase in the degree of synchrony is also evident in the average activity time courses in panel B. Increasing interCC is therefore a very effective way of increasing the number of CEs that are part of SEs. As network synchronization increased with higher interCC values, the number of CEs also increased, as illustrated in the left panel of Fig. 5C. In addition, the standard deviation in the number of CEs among the clusters decreased with an increase in interCC, again indicating that the cluster activity became more uniform when interCC was increased. Finally, the fraction of synchronization events in which all 5 clusters participated (frequency of NCE_5_) increases with the interCC (right panel of Fig. 5C), again demonstrating the tendency of intercluster coupling to increase the degree of synchronization between clusters.

These simulations of the effect of increasing intercluster coupling provide results similar to those described in [22] in which tachykinin receptor antagonists were used to block the effects of NKB. NKB has excitatory effects, acting on multiple tachykinin receptors [34], so blocking its action will reduce the effective coupling among neurons. In the brain slice studies, blocking the tachykinin receptors reduced the number of events, while in vivo blocking the receptors reduced the average amplitude of the SEs. These effects are consistent with reducing the interCC in our simulations: the number of CEs declines as does the number of clusters participating in the SEs (Fig. 5).

We next followed the same procedure, but this time keeping the intercluster coupling parameter fixed and varying the intracluster coupling. Figure 6A shows histograms of the number of clusters participating in events during 40 s simulations, with intraCC from 60% (top) to 95% (bottom). As intraCC is increased, there are more events with some level of synchronized activity. However, for all values of the parameter investigated, there were more mSEs than SEs. Panel B shows average activity time courses over 5 s for three cases (tan shading in the histograms) that show that while synchronous events occur, there are many instances in which single clusters fire alone. Increasing intraCC does increase the number of CEs, but the standard deviation across the clusters changes little (Fig 6C, left), again indicating that increasing intraCC is not particularly effective at bringing all clusters into synchrony. Interestingly, increasing the intraCC led to a reduction in the fraction of SEs in which all 5 clusters participated (Fig 6C, right). That is, increasing the intracluster coupling lowers the degree of synchrony among the SEs.

**Fig 6.**
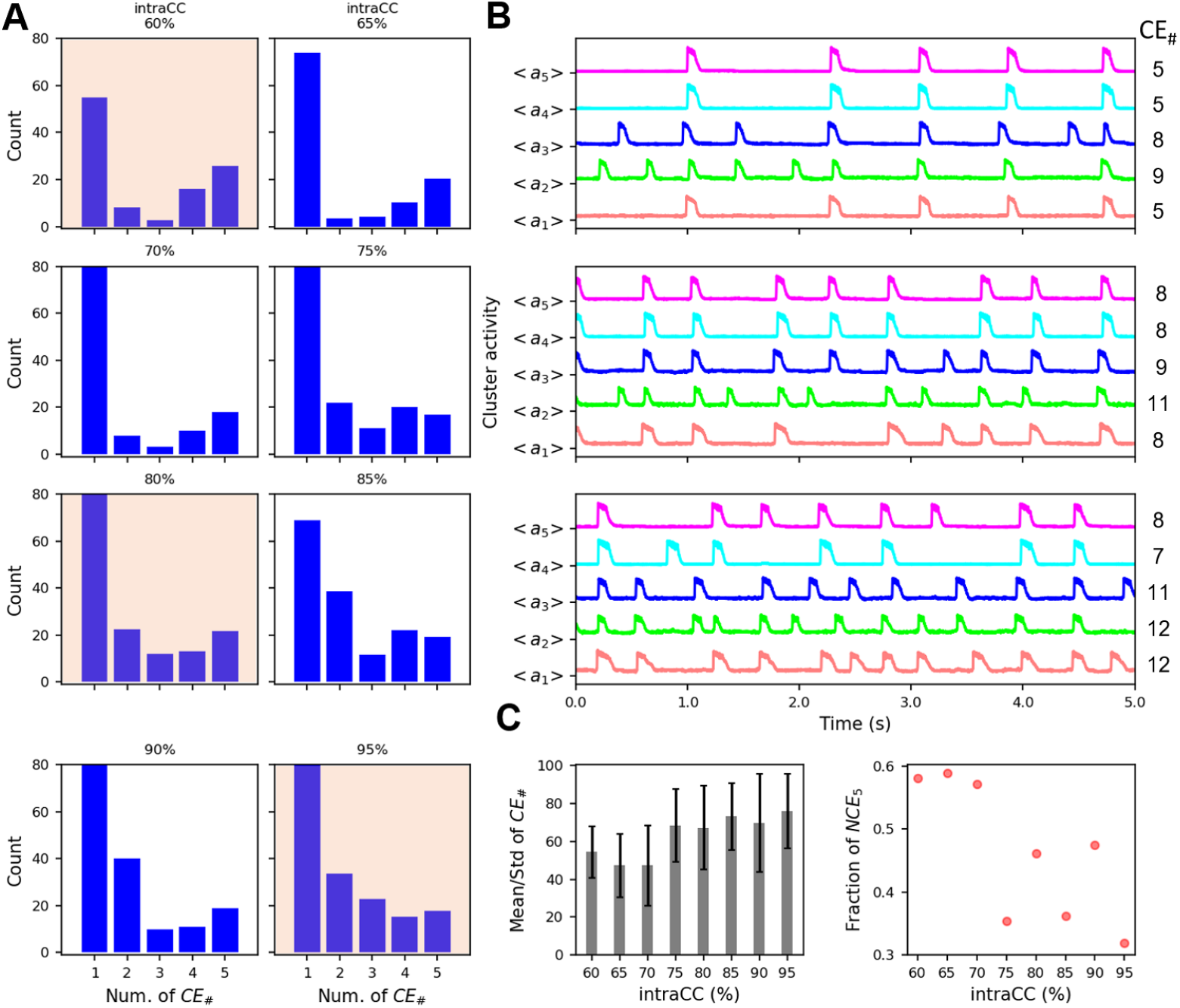
Increasing intracluster coupling has a weak effect on the number of cluster events, and weakly decreases cluster synchronization during SEs. A: Histograms showing the number of events in which 1, 2, …, 5 clusters fire together during simulations with 40 s duration. The intraCC is increased moving from top to bottom. B: Average activity time courses of the 5 clusters during 5 s of simulation time, corresponding to the histograms with tan shading in the previous panel. C: (left) Mean number of CEs, along with standard deviation, for the range of intraCC values explored in the histograms. (right) The fraction of SEs in which all five clusters participate, NCE_5_/NSE. InterCC=0.2%.

This paradoxical finding, that the mean size of SEs would be lower with higher levels of intracluster coupling, mirrors a similar paradoxical finding reported in [22]. Here, it was reported that blocking Dyn receptors with the opioid receptor antagonist norBNI had inconclusive effects on the number of events in brain slice studies of KNDy neurons, but that in vivo the antagonist significantly reduced the amplitude of SEs by 51%. Since Dyn is an inhibitory neurotransmitter, this means that inhibiting an inhibitor decreased the size of the SEs. This finding is in agreement with results from the modular model if one postulates that Dyn acts primarily to inhibit intracluster coupling. If so, then Dyn would lower the intraCC, increasing NCE_5_/NSE and thereby increasing the mean size of the SEs. Blocking its effect would increase intraCC, decreasing NCE_5_/NSE and thus decreasing the mean SE size as reported in [22].

## Discussion

This study was motivated by recent data describing neuronal activity within the arcuate nucleus KNDy neuron network in vivo, at single-cell resolution [22, 26]. The data described how single neurons coordinate to generate the synchronized network events that drive pulsatile LH release. It led Han et al (2023) to propose a new paradigm for the synchronization events, where glutamate transmission provides the main synchronization drive, and Dyn and NKB play supporting roles, amplifying the synchronization. The single-cell resolution also enabled the observation of leader cells that activate first during synchronization events, with a consistent order of recruitment over synchronization events [26]. At the same time, some networks exhibited more variability in the order of firing. We have demonstrated here that these experimental findings can be explained by a network of neurons that has a modular structure consisting of clusters of highly connected neurons with sparse intercluster coupling.

We provide an explanation for why temporal ordering could be seen as fixed or variable in different studies, or different animals. The findings of Moore et al. suggest that there are distinct leader and follower cell populations [26]. The consistency of firing order between groups of cells suggests a cluster organization in the KNDy network, like the one we have adopted. This experimental study, and our simulations that exhibit a high correlation between cell ID and firing order, show “blockiness” in the ID versus order scatter plot (Fig 4Bii in [26] and our Fig 3A). This suggests that the same groups of cells are consistently activated around the same time relative to the start of a synchronization event. In our simulations, this is due to the sequential recruitment of different clusters to an active state. This happens when intercluster connectivity is high and intracluster connectivity is low. In other words, we get a consistent order of firing between cells when the coupling between clusters is sufficiently strong that the most active cluster can consistently evoke episodes of activity in less-active clusters, establishing a leader-follower hierarchy. Also, the coupling within a cluster is sufficiently weak so that while some clusters are active even without intercluster coupling, other clusters are silent without the influence of other clusters. Paradoxically, this reveals the cluster organization.

On the other hand, when the network is more modular (high intracluster connectivity and low intercluster connectivity), the consistency in the order of cell activations is lower. This is because the clusters are more independent and any cluster can generate a CE that then may trigger a more global SE. Because each cluster is as likely as another to trigger a synchronization event, the ordering of cell activations is variable. This more variable temporal ordering is more consistent with results from Han et al. [22]. We point out, however, that relatively small connectivity parameter manipulations were sufficient to go from consistent to variable temporal ordering, suggesting that minor connectivity differences in the experiments from Han et al. versus Moore et al. can explain the difference in their results. Such differences could reflect the sex difference between animals used in the experiments. Male mice were used in Han et al. and ovariectomized female mice were used in Moore et al. In a subsequent study of KNDy neurons in brain slices from female mice, there was little consistency in the order of spiking during mSEs [23], consistent with the findings from male mice by the same lab [22]. In addition to sex differences, the variation in connectivity parameters could also be due to different levels of Dyn and NKB released from KNDy neurons, or differences in receptor expression for these neurotransmitters.

One paradoxical finding reported in [22] was that blocking receptors for either NKB or Dyn reduced the size of the SEs, indicating that these neurotransmitters both act to increase synchrony among KNDy neurons, even though one is excitatory and the other inhibitory. Our simulations indicate that this could be explained if the KNDy neuron network is modular and if NKB acts primarily to increase intercluster coupling and Dyn acts primarily to decrease intracluster coupling. This could occur if there is a heterogeneous distribution of receptors for the two neurotransmitters. Indeed, in male mice it was found that less than half of the KNDy neurons express the *κ*-opioid receptors for Dyn [35]. Also, Kiss, Dyn, and NKB are packaged into separate vesicles [36], so the proportion of neurotransmitter type released at synapses likely varies from synapse to synapse.

One notable difference between the SEs produced by our model network and those observed in actual KNDy neuron populations [22, 26] is the much shorter time between SEs and SE duration in the simulations. Replicating the much slower SEs reported in the experiments would greatly increase the time required for computer simulations, without changing the results of interest to us, such as the order of spiking during SEs and participation of neurons in some, but not all SEs. Indeed, as reported in [22], the SEs recorded in brain slices had much shorter inter-SE intervals than those recorded in vivo, but the basic properties of the SEs were the same.

The simulations were performed with a version of the Hodgkin-Huxley model, with only two voltage-dependent ion channel types. The actual KNDy neurons are almost certainly more complicated, but to date no biophysical model of KNDy neurons based on single-cell data has been published. We do not believe, however, that the use of more complete single-cell models would impact the findings of this study, which are determined primarily by the network structure.

A more fundamental assumption that we made is that the coupling between neurons is through glutamate, and that NKB and Dyn act as modulators of that coupling. This is consistent with recent results from [22], but is contrary to the proposal that the episodes of activity are started by the actions of NKB and terminated by the actions of Dyn [17]. In our model, the excitatory action of glutamate declines over time due to synaptic depression, which is responsible for terminating each activity episode, as in developing networks [27]. Another possibility is that the buildup of some intrinsic hyperpolarizing current or currents could cause episode termination [27]. Indeed, [22] found evidence for Ca^2+^-activated K^+^ current in KNDy neurons that could play such a role. The presence of these currents does not, however, discount the potential role of synaptic depression in episode termination.

The key property of the networks used in our study is that they are modular, consisting of clusters of highly-coupled neurons with sparse coupling between clusters. Our key findings cannot be replicated in a non-modular network. When looking at a homogeneous network (i.e. a single cluster), we found that all neurons consistently participate, or not, in synchronization events. This is contrary to the finding that KNDy neurons participate in some, but not all, SEs [22]. We also found that the temporal order of spiking during a CE is similar from one event to the other; there are definite leader cells and follower cells (Fig 4C). This order is set by the background current in each cell, i.e., their level of excitability. Thus, with a single cluster we do not capture the variable order of spiking during SEs reported by [22]. If noise were included in simulations, then this variability would likely occur, but then it would be difficult to capture the high degree of temporal ordering reported in [26] and, in some cases, in [22]. Finally, with a single cluster we would have no explanation for how both NKB and Dyn could increase the size of SEs, since our explanation relied on having two independent coupling parameters, with NKB acting primarily on one and Dyn primarily on the other.

The modular network hypothesis put forward here makes two experimentally testable predictions. First, application of Dyn or NKB, in addition to strengthening SEs, should change the recruitment order variability as shown in Fig 4A. Second, in homogeneous networks the order of recruitment is mostly determined by cell excitability: the more excitable cells fire before the least excitable cells. In the modular network, the cells that consistently fire first are the ones that belong to the most excitable clusters. So recruitment order does not depend on cell excitability, it depends on cluster identity. That is, for neurons in a modular network, it is not “who they are” that determines recruitment order, but “who they know”. Thus we predict that in networks with consistent recruitment order, the cells that are recruited first are not necessarily the most excitable cells.

## Conclusion

A mathematical model of the KNDy network with a modular structure elegantly explains the features of KNDy population activity observed experimentally:

1. Different SEs involve different combinations of clusters – thus cells participate in SEs according to the cluster they belong to.

2. The balance between intra- and inter-cluster connectivity determines whether the recruitment order of cells during SEs is consistent or not.

When recruitment is consistent across SEs, this is because the order of cluster recruitment is consistent.

3. The actions of the excitatory modulator NKB and the inhibitory modulator Dyn both lead to a greater degree of synchronization during SEs if they act on distinct sets (intercluster versus intracluster) of synaptic connections.

## Acknowledgments

W.B. thanks the Department of Mathematics and the Institute of Molecular Biophysics at Florida State University for hosting his sabbatical.

## Notes

### Competing Interest Statement

The authors have declared no competing interest.

